# Sirtuin gene isoforms and genomic sequences of mouse and humans: Divergence across species

**DOI:** 10.1101/2022.04.18.488659

**Authors:** Xiaomin Zhang, Gohar Azhar, Fathima S. Ameer, Jasmine Crane, Jeanne Y. Wei

## Abstract

**Background:** There are seven sirtuin genes in the mammalian genome. Each sirtuin gene contains multiple exons, and is likely to undergo alternative splicing, thereby increasing sirtuin gene diversity. Since the alternatively spliced isoforms tend to increase with advancing age, it is important to study the effect of sequence change on isoform function. Additionally, the divergence of isoform patterns between human and mouse will help us to properly interpret the findings from animal models, especially in age-related studies. Recently, more than 20 human sirtuin isoforms have been identified, but whether the mouse genome might have similar isoforms remains incompletely established.

**Methods:** The mRNA, protein and genomic DNA sequences of mouse sirtuin genes, as well as the transcription factor binding sites, including that of SRF, were analyzed. A cellular stress model with serum deprivation and restoration was used to reflect the blood supply and nutrients level changes in the ischemia and reperfusion condition, and the expression of sirtuin isoform was assessed.

**Results:** Here, we report the identification of 15 mouse sirtuin isoforms, of which over half have not been previously reported. Exon skipping was the main event, which led to domain losses in the nuclear localization signal, nucleolar localization signal, and/or the mitochondrial targeting sequence among sirtuin isoforms. Among 7 sirtuin genes, 6 mouse sirtuin genes had different exon numbers versus that of human sirtuin genes. Only the sirtuin-2 gene had the same number of exons in both human and mouse. However, there were differences in the sirtuin-2 gene isoforms and the regulatory domains between the two species. The expression of sirtuin gene isoforms under serum stress was also different.

**Conclusions:** Alternative splicing increases both sirtuin transcriptome and proteome diversity. However, the sirtuin isoforms were not well conserved between human and mouse, which should be taken into consideration when extrapolating animal studies for human physiology and pathology. Our results will help to elucidate the role of sirtuin genes in the regulation of cellular stress response, including ischemia and reperfusion. We propose that the existence of the CArG and CArG-like sequences in sirtuin genes may imply a role for SRF in the sirtuin family transcriptional regulation.

## Introduction

Alternative splicing (AS) is the RNA process by which various selections of splice sites in precursor mRNA (pre-mRNA) result in more than one distinct mRNA and protein variants [1–4]. It has been estimated that most of the multi-exon genes are alternatively spliced, and many splice variants are differentially expressed in a tissue-specific manner [5,6].

The sirtuin proteins belong to the class III histone deacetylases (HDACs), which regulate histones and other non-histone proteins via reversible lysine acetylation. To date, seven homologs have been identified (SIRT1 to SIRT7) in the mammalian genome [7], all of which contain multiple exons, and are likely to undergo alternative splicing. The pre-mRNA splicing process is mainly catalyzed by the spliceosome, a multi-mega dalton ribonucleoprotein (RNP) complex, but it can also be influenced by the other proteins, including RNA binding proteins and transcription factors [8]. Environmental stresses, such as the deprivation of growth factors, may also influence the outcome of the alternative splicing event [9].

More than 20 human sirtuin isoforms have been identified, but whether the mouse genome might have similar isoforms is unknown [10]. Since the mouse models have been extensively used in biological studies, and the results are being used to translate into human physiology, it is worthwhile to understand the potential impact of isoform divergence on transcriptome and/or proteome function between human and mouse. In addition, alternatvely spliced isoforms tend to increase with advancing age, thus it is important to study the “repertoire” of sirtuin isoforms in the mouse versus human.

In the present report, we identified 15 mouse sirtuin isoforms, over half of which have not been previously reported, and the genomic landscape of the sirtuin genes in mouse was compared with that of human. Furthermore, the transcription factor binding sequences, including the serum response element sequences, were analyzed.

## Materails and methods

### Nomenclature of the sirtuin isoforms and bioinformatics analysis

All the mRNA, protein and genomic DNA sequences of the sirtuin genes were obtained from The National Center for Biotechnology Information (NCBI) Reference Sequence (RefSeq) database [11]. The nomenclature of the sirtuin isoforms follows the guidelines used in The RefSeq database, in which the constitutively spliced form of sirtuin gene is termed as isoform-1, and alternatively spliced isoforms are named as isoform-2, isoform-3, and so on [11]. For instance, the SIRT1 isoform-1 (termed SIRT1v1, accession number NM_019812 for mRNA, and NP_062786 for protein) represented the sirtuin-1 gene (the homolog of the Sir2 protein), which had been widely studied for many years [12], the second SIRT1 isoform was then named SIRT1v2 (accession number NM_001159589 for mRNA, and NP_001153061 for protein). The data collection for this report was completed as of February 21, 2021.

### Bioinformatics analysis

The web-based software “Blast” and “MultAlin” were used for sequence alignment [13]; the “ScanProsite” software was used to determine the conserved domain of the sirtuin isoform [14]; the SWISS-MODEL software was used for analysis of the 3-D protein structure [15]. The software NLS Mapper [16], NLSdb [17], NetNES [18], LocNES [19], and NucPred [20] were used for the analysis of nuclear localization signal (NLS) and the nuclear exporting signal (NES) sequences. The nucleolar localization signal (NoLs) sequence was analyzed using NOD (Nucleolar Localization Sequence Detector) [20]. The software MITOPROTII [21], TargetP [22], and MitoFates [23] were employed for the analysis of mitochondrial targeting sequence (MTS).

### Cell culture and a cellular stress model

The DMEM, newborn bovine serum (NBS), lipofetamin 2000 were purchased from Thermo Fisher Scientific (Waltham, MA). C2C12 cell line (ATCC CRL-1772) was obtained from ATCC. The C2C12 cells were maintained in DMEM containing 10% NBS. The cellular stress model with serum deprivation and restoration was similar to what was described previously [24]. For serum deprivation treatment, the cells were washed with PBS twice and then incubated in DMEM containing 0.2 % serum for 3, 6, 24, and 48 hours as indicated. For serum deprivation and restoration treatment, the cells were first subjected to serum deprivation for 48 hours and then were cultured in DMEM containing 10% NBS for an additional 6, 18, 24, and 48 hours, respectively [24].

### RNA isolation and Quantitative Reverse-transcriptase PCR (qRT-PCR)

All RNA samples were isolated from mouse tissues using the miRNeasy Mini kit and RNase-free DNase I (Qiagen) [25]. The reverse transcription and RT-PCR reagents were obtained from Applied Biosystems. The isoform specific primers were selected from the unique sequence region of each isoform sequence using Primer-Blast (NCBI) and synthesized by Integrated DNA Technologies (Coralville, Iowa). The primers for the detection of the mouse sirtuin isoforms were as follows: SIRT1v1 forward ttgaccgatggactcctcac, reverse gtcactagagctggcgtgtg. SIRT1v2 forward cggctaccgaggtccatatac, reverse agctcaggtggaggaattgt. SIRT2v1 forward gcaggaggctcaggattcag, reverse agacgctccttttgggaacc. SIRT2v2 forward gagccggaccgattcagac, reverse agacgctccttttgggaacc. SIRT2v3 forward tttggtgggagccggaatc, reverse gcctctgggtaaggaaggtg. SIRT3 forward tcacaaccccaagccctttt, reverse agtgagtgacattgggcctg. SIRT4 forward tctcctctcaccaacccaac, reverse tccacgttctgagtcaccaa. SIRT5 forward gccaccgacagattcaggt, reverse tttccacagggcggttaaga. SIRT6 forward cattgtcaacctgcaaccca, reverse cctgcttcatgagtctgcac. SIRT7 forward tctacaaccggtggcaggat, reverse catagtgacttcctactgtggct. The 5S RNA reference primers were used as endogenous control [25].

The PCR amplification was performed in a StepOnePlus Real-Time PCR System (Applied Biosystems). Relative expression values were obtained by normalizing CT values of the mRNA genes in comparison with CT values of the endogenous control (5S RNA) using the CT method [25,26].

### Statistical analysis

Data were given as mean values ± SD, with n denoting the number of experiments unless otherwise indicated. A two-tailed T-test was used to determine the differences between the two groups. A p value of < 0.05 was considered to be statistically significant.

## Results

### The sirtuin isoforms in the mouse genome

There were 15 annotated mouse sirtuin isoforms in the NCBI RefSeq Database, which were derived from the seven sirtuin genes (SIRT1-SIRT7) that are located at different gene loci on different chromosomes (Table 1). Six of the seven sirtuin genes, excluding SIRT5, had more than one isoform. SIRT1, SIRT4, SIRT6, and SIRT7 had two isoforms, whereas SIRT2 and SIRT3 each had 3 isoforms (Table 1 and Fig 1). The nomenclature of the isoforms follows what was used in the NCBI RefSeq database as described in “Methods and Materials”.

**Table 1.** Mouse sirtuin genes and their isoforms. Alternative splicing resulted in the loss of part of the sequence and changed the length of most of the alternatively spliced isoforms.

| Gene | Chromosome | Band/Region | Number of Exon | DNA | Isoform | mRNA | Protein | Amino Acid | Sirtuin domain |
| --- | --- | --- | --- | --- | --- | --- | --- | --- | --- |
| SIRT1 | 10 | 10 B4 | 9 | NC_000076.7 (63154784..63174814, complement) | V1 | NM_019812.3 | NP_062786 | 737 | 246-481 |
|  |  |  |  |  | V2 | NM_001159589.2 | NP_001153061.1 | 698 | 207-442 |
| SIRT2 | 7 | 7 B1 | 17 | NC_000073.7 (28466177..28488090) | V1 | NM_022432.4 | NP_071877.3 | 389 | 77-331 |
|  |  |  |  |  | V2 | NM_001122765.1 | NP_001116237 | 352 | 40-294 |
|  |  |  |  |  | V3 | NM_001122766.1 | NP_001116238.1 | 319 | 7-261 |
| SIRT3 | 7 | 7 F4-F5 | 8 | NC_000073.7 (140443576..140462222, complement) | V1 | NM_022433.2 | NP_071878.2 | 257 | 1-231 |
|  |  |  |  |  | V2 | NM_001127351.1 | NP_001120823.1 | 257 | 1-231 |
|  |  |  |  |  | V3 | NM_001177804.1 | NP_001171275.1 | 334 | 74-308 |
| SIRT4 | 5 | 5F | 7 | NC_000071.7 (115616065..115628322, complement) | V1 | NM_001167691.1 | NP_001161163.1 | 333 | 44-305 |
|  |  |  |  |  | V2 | NM_133760.1 | NP_598521.1 | 333 | 44-305 |
| SIRT5 | 13 | 13 A4 | 13 | NC_000079.7 (43518849..43549073) |  | NM_178848.3 | NP_849179.1 | 310 | 51-301 |
| SIRT6 | 10 | 10 C1 | 11 | NC_000076.7 (81457621..81463631, complement) | V1 | NM_181586.4 | NP_853617.1 | 334 | 45-257 |
|  |  |  |  |  | V2 | NM_001163430.2 | NP_001156902.1 | 294 | 5-217 |
| SIRT7 | 11 | 11 E2 | 12 | NC_000077.7 (120509197..120515840, complement) | V1 | NM_153056.3 | NP_694696.2 | 402 | 102-315 |
|  |  |  |  |  | V2 | NM_001363439.1 | NP_001350368.1 | 382 | 102-315 |

**Fig 1.**
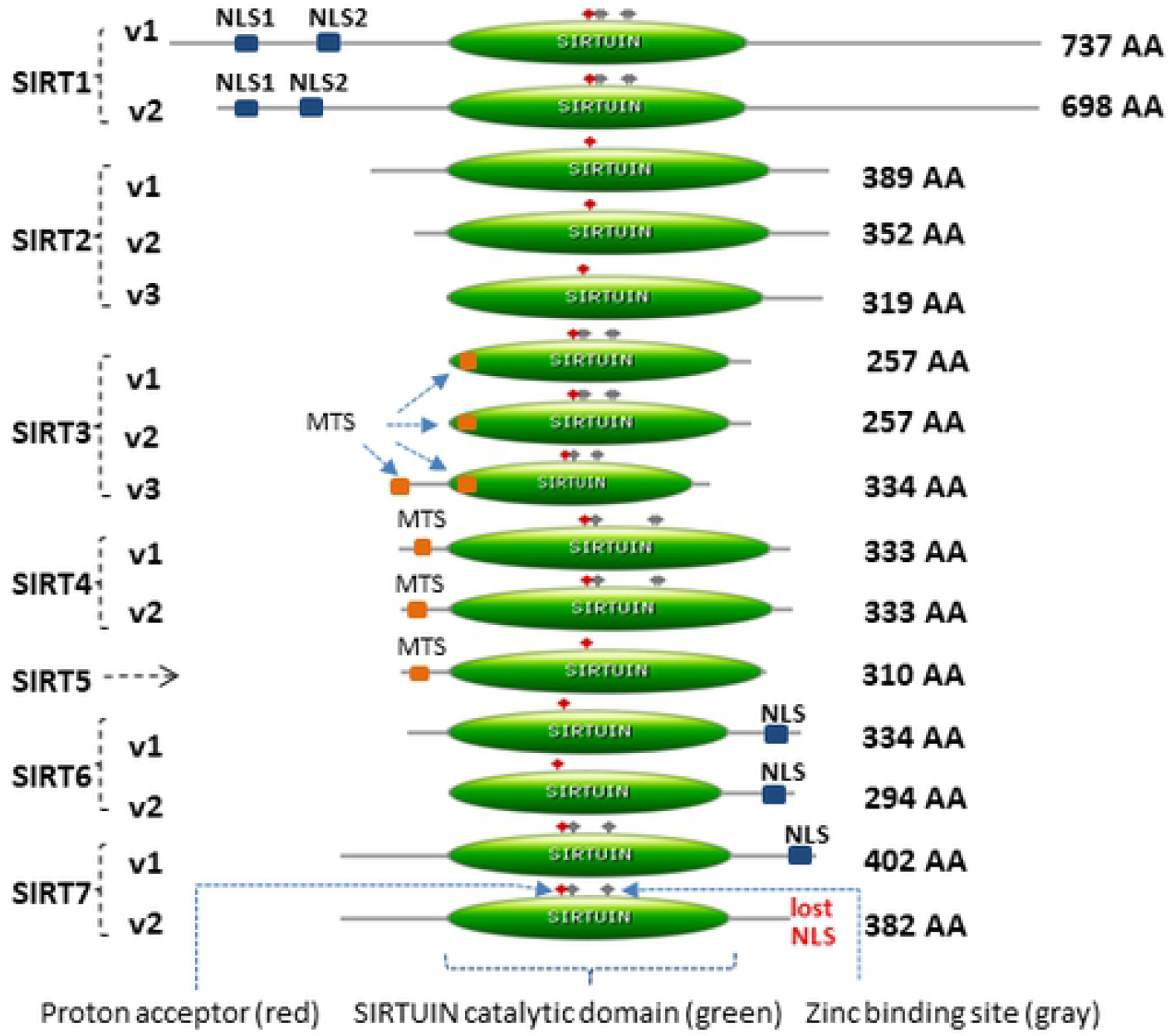
The schematic diagram of 15 isoforms of mouse sirtuin genes (SIRT1 - SIRT7). The loss of part of the protein domain was observed among isoforms, which included part of the N- or C-terminus, catalytic domain (in green color), nuclear localization signal (NLS, in navy color), and mitochondrial targeting signal (MTS, in orange color). The functional domain analysis was performed using the “ScanProsite” software (please see Methods). AA refers to amino acid.

The SIRT1 gene had two isoforms, SIRT1v1 represents the longer transcript and encodes the longer isoform, the SIRT1v2 lacked an alternate in-frame exon (exon-2) in the 5’ coding region compared to variant 1, and therefore the v2 isoform was shorter than isoform v1.

The SIRT2 had three isoforms. The SIRT2v1 represented the longest transcript and encoded the longest isoform. Compared to SIRT2v1, the v2 isoform lacked an alternate in-frame exon in the 5’ coding region and used a downstream start codon, thus the v2 protein was 37 amino acides shorter at the N-terminus. Compared to SIRT2v1, the v3 isoform lacked multiple in-frame exons (ex2, 3, 4) in the 5’ coding region, resulting in an even shorter N-terminus in the SIRT2v3 isoform protein.

SIRT3 had three isoforms. The isoforms v1 and v2 had slightly different lengths of mRNA transcripts, but encoded the same length of proteins, both of which were 257 amino acides in length. The SIRT3v3 was the longest one among the three isoforms, which was 334 amino acids in length that was almost 78 amino acids longer than the v1 and v2 proteins.

SIRT4 had two mRNA transcripts, v1 was 1645 bp in length whereas v2 was 1553 bp in length, but both transcripts encoded the identical protein isoforms with the same length of 333 amino acids.

SIRT5 had only one isoform with a transcript of 1369 bp that encoded a protein of 310 amino acides in length.

SIRT6 had two isoforms, v1 was 334 amino acides in length, and v2 was 294 amino acids in length.

SIRT7 had two isoforms, isoform v1 was 402 amino acides in length and v2 was 382 amino acids in length.

Analysis of the sirtuin isoforms and exon sequences revealed that phenomenon of exon skipping happened in majority of the alternatively spliced sirtuin isoforms. The length of the isoform proteins ranged from 257 amino acids (e.g., SIRT3v3) to 737 amino acids (e.g., SIRT1v1) (Table 1).

To determine the functional domain of all the sirtuin isoforms, the protein sequences of 15 isoforms were analyzed using the web-based software “ScanProsite” [14]. As shown in Table 1 and Fig 1, the sirtuin catalytic domain length ranged from 212 amino acids (e.g., SIRT6v2) to 261 amino acids (e.g., SIRT4v1). Exon skipping caused the alternatively spliced isoforms to lose part of the protein domain.

To examine the potential structural changes due to a domain loss, all 15 sirtuin isoforms were subjected to protein structure homology-modelling using the SWISS-MODEL software [15]. As shown in Fig 2, protein domain loss resulted in minor to major structural changes among the alternatively spliced isoforms derived from the same sirtuin gene.

**Fig 2.**
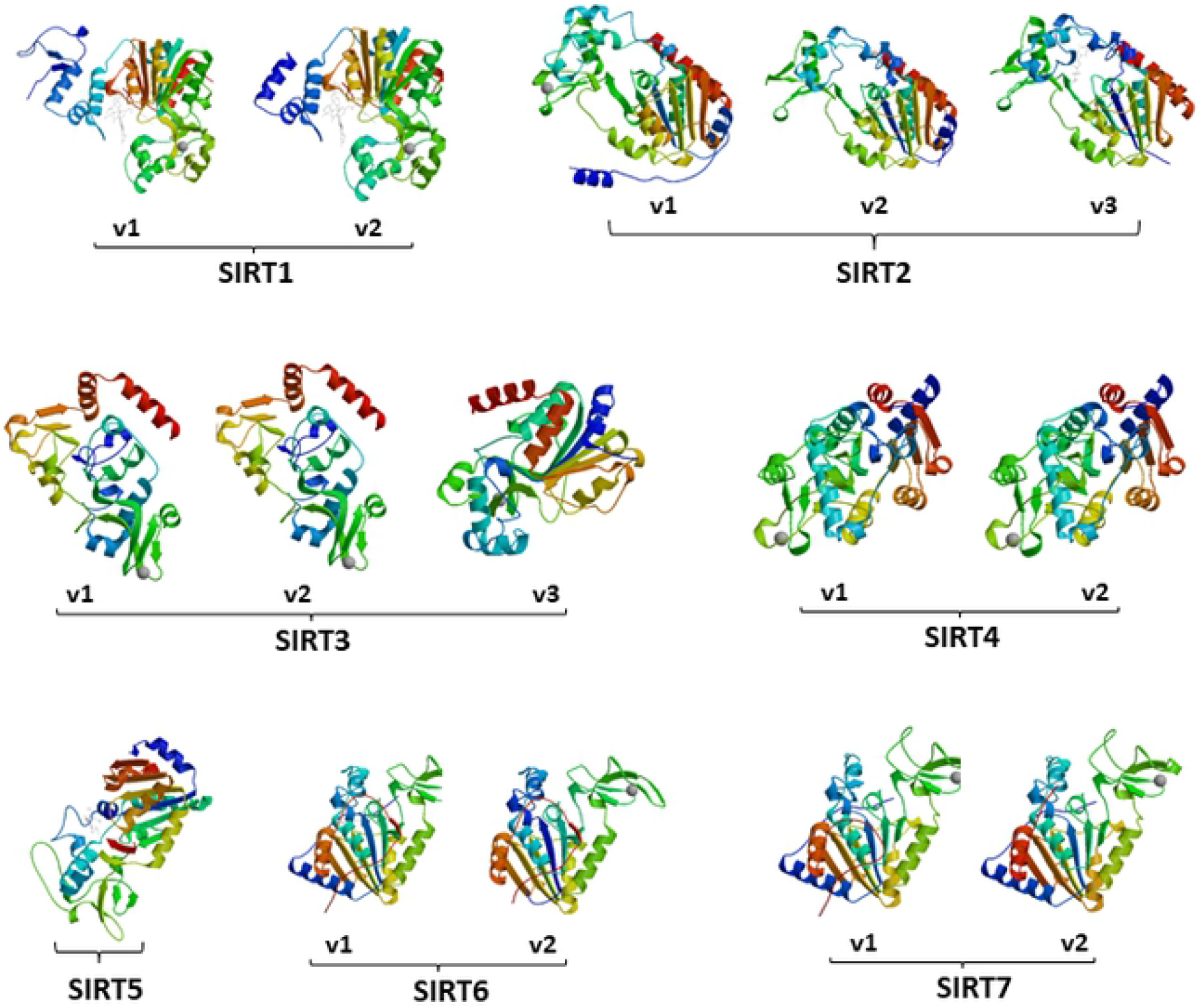
The 3-D protein structure of mouse sirtuin isoforms that were generated using SWISS-MODEL software.

### Analysis of subcellular localization sequences of sirtuin isoforms

The sirtuin isoform protein sequences were subjected to an evaluation for the presence of nuclear localization signal, nucleolus localization signal, and the mitochondrial targeting sequence signal.

Three sirtuin genes, SIRT1, SIRT6 and SIRT7, each contained the exon with nuclear localization signal. The preservation of the exon with NLS predicts the isoform’s capability of nuclear localization.

SIRT1v1 contained two NLS sequences, NLS1 (PLRKRPRR) was in exon-1, and NLS2 (PPKRKKRK) was in exon-3. Since SIRT1v2 skipped exon-2, but retained exon-1 and exon-3, it therefore contained both of the two NLS sequences of SIRT1v1 as well (Fig 3A). Both v1 and v2 isoforms were predicted to be localized in the nucleus.

**Fig 3.**
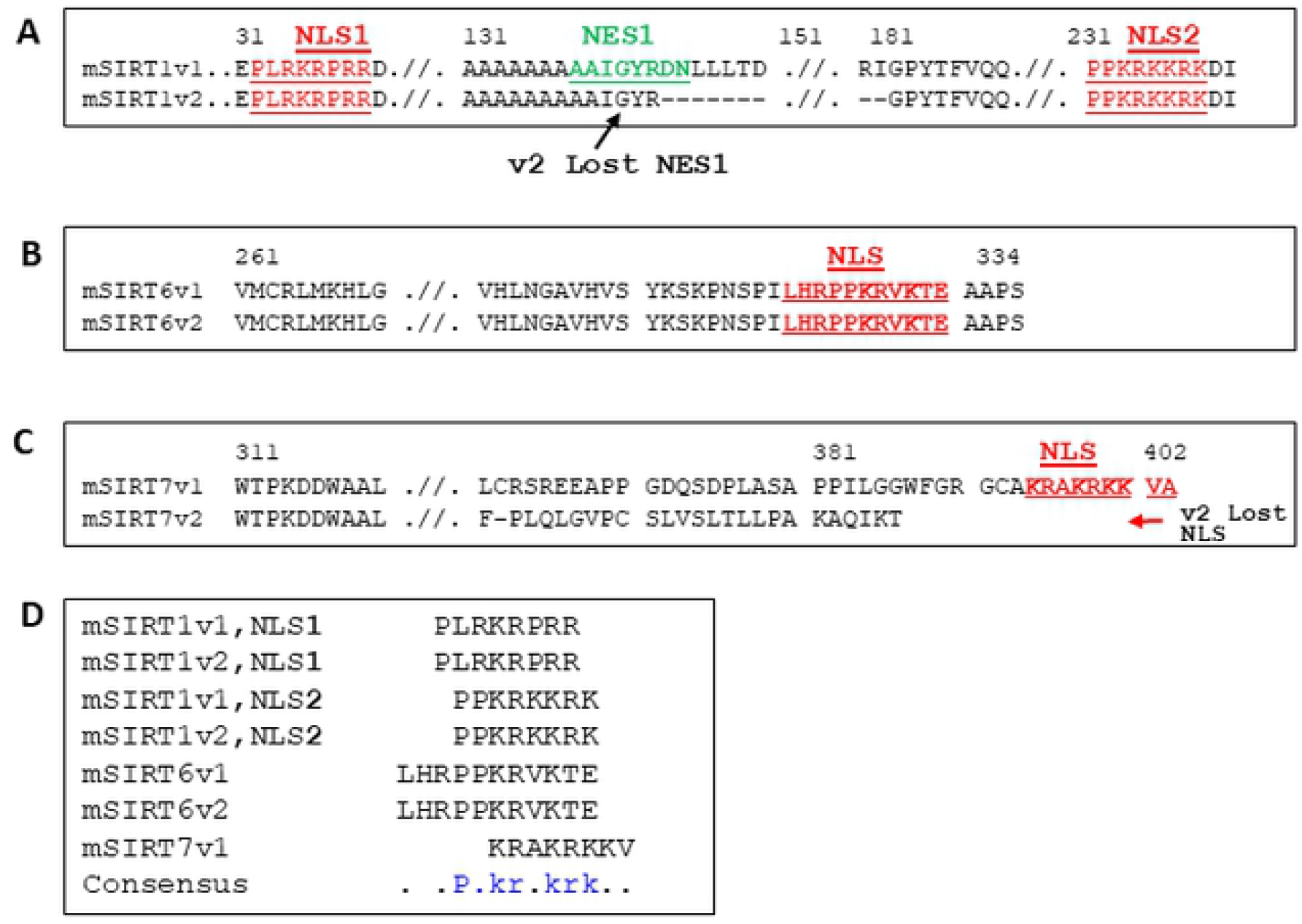
The nuclear localization signal (NLS) sequences in mouse sirtuin isoforms derived form mouse SIRT1, SIRT6 and SIRT7 genes. 3A. Both SIRT1v and SIRT1v2 had two NLSs (NLS1 and NLS2), but SIRT1v2 lost NES1 (nuclear export signal). 3B. Domain loss happened to the N-terminus in SIRT6v2 (sequence not shown), and did not affect the NLS region. Therefore, both SIRT6v1 and SIRT6v2 had NLSs. 3C. SIRT7v2 lost its NLS. 3D. The NLS consensus sequences among the SIRT1, SIRT6 and SIRT7 isoforms.

SIRT6v1 had one NLS sequence (LHRPPKRVKTE) at the COOH-terminus of the protein. The SIRT6v2 isoform had an exon skipped at the NH-terminus, but retained the NLS (LHRPPKRVKTE) sequence at the COOH-terminus. The Mouse SIRT6 v2 aparently lost 40 AA at the N-terminus, which was not a part of the catalytic domain; the rest of the protein sequence between v1 and v2 remained the same. Therefore, v2 retained the NLS (Fig 3B).

SIRT7v1 had a NLS sequence (KRAKRKKV) at the COOH-terminus, whereas SIRT7v2 skipped the exon containing the NLS sequence (Fig 3C).

Alignment of the NLS sequences of the five isoform proteins of SIRT1, SIRT6, and SIRT7 revealed that the isoforms all shared a consensus sequence “P·KX·KRK” (Fig 3D).

The existence of the nucleolar localization sequences (NoLSs) was also examined in the sirtuin isoforms with the software (http://www.compbio.dundee.ac.uk/www-nod/HelpDoc.jsp) [27]. Both SIRT1v1 and SIRT1v2 were found to have two NoLS sequences. One NoLS was “ADEPLRKRPRRDGPGLGRSPG” at N-terminus (between positions 28 and 48 in both v1 and v2), and the other NoLS “VINILSEPPKRKKRKDINTIEDAV” was at C-terminus (between positions 216 and 239 in SIRT1v1, and between positions 177 and 200 in SIRT1v2) (Fig 4A, 4B).

**Fig 4.**
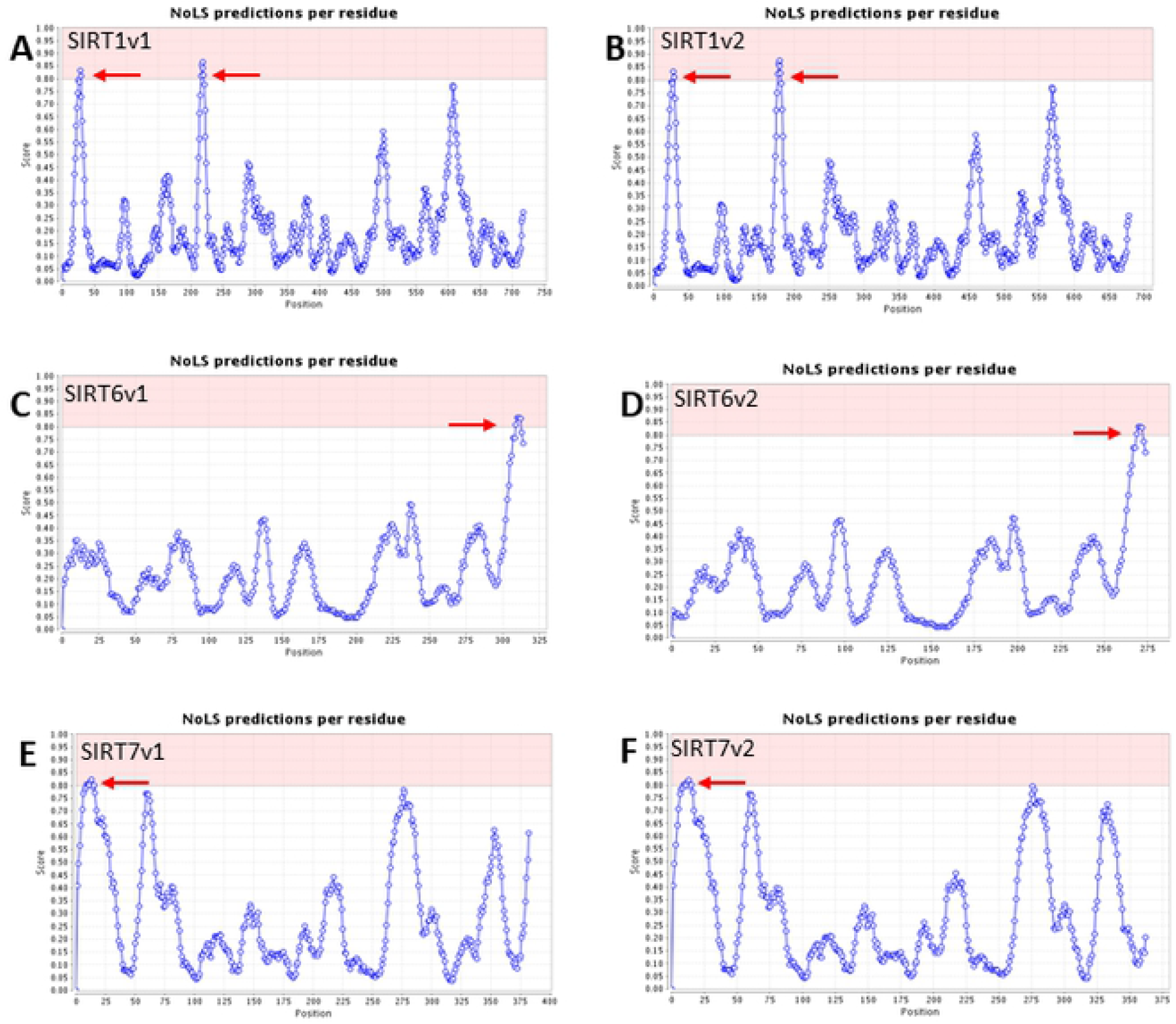
Analysis of nucleolar localization signal (NoLS) sequences in the mouse sirtuin isoforms. The red arrow pointed to the predicted NoLS sequence. A. Two NoLSs were detected at the N-terminal region of SIRT1v1 isoforms. B. Two NoLSs were detected at the N-terminal region of SIRT1v2 isoforms. C. One NoLS was detected at the C-terminal region of SIRT6v1 isoform. D. One NoLS was detected at the C-terminal region of SIRT6v2 isoform. E. One NoLS was detected at the N-terminal region of SIRT7v1 isoform. F. One NoLS was detected at the N-terminal region of SIRT7v2 isoform.

Both SIRT6v1 and SIRT6v2 had one NoLS “VSYKSKPNSPILHRPPKRVKTEA” at the C-terminus (between positions 309 and 331 in SIRT6v1, and between positions 269 and 291 in SIRT6v2, respectively) (Figs 4C and 4D).

Both SIRT7v1 and SIRT1v2 had one NoLS “RSERKAAERVRRLREEQQRERLRQVS” at the N-terminus (between positions 9 and 34 in both SIRT7v1 and v2) (Figs 4E and 4F).

Software tools that can detect the mitochondrial targeting sequence (MTS) were employed to examine the existence of MTS in the sirtuin isoform proteins. They included MITOPROTII [21], TargetP [22] and MitoFates [23].

The SIRT3 gene had three isoforms. SIRT3v1 and v2 isoforms both encoded the identical protein, which had a MTS sequence “VTHYFLRLLHDKELLLRLYTQNI”. The SIRT3v3 isoform was the longest isoform among the three isoform; it had a MTS sequence, “VTHYFLRLLHDKELLLRLYTQNI”, which was the same as that of v1 and v2. In addition, the v3 had another MTS sequence “MALDPL GAVVLQSIMA LSGRLALAAL RLWGPGGGRR PISLCVGASG” (Fig 5A).

**Fig 5.**
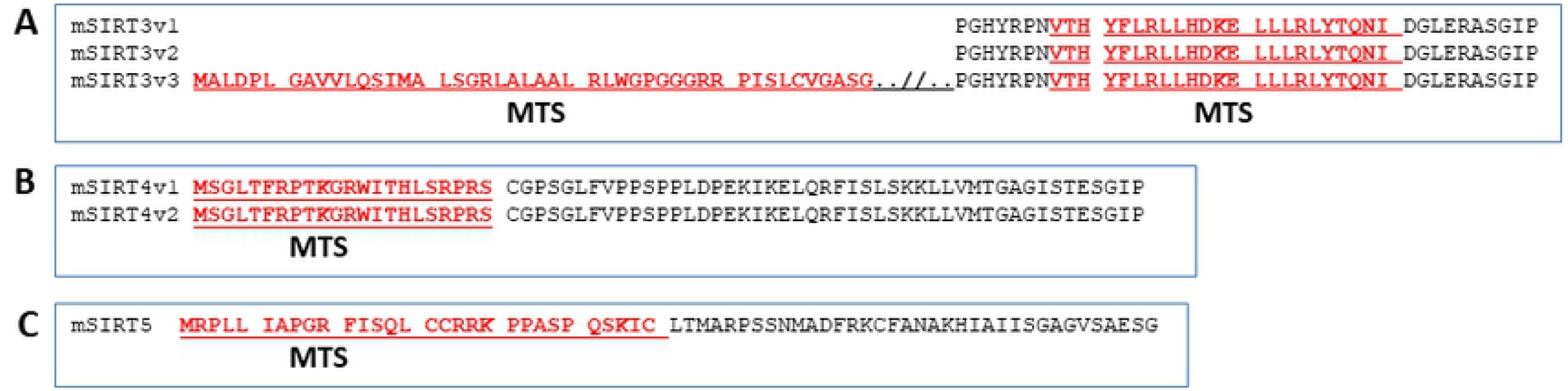
Mitochondrial targeting sequence (MTS) in mouse SIRT3 (A), SIRT4 (B) and SIRT5 (C) isoforms.

The SIRT4 gene had two mRNA isoforms with slightly different mRNA lengths but both encoded the same isoform protein, which had a MTS sequence “MSGLTFRPTKGRWITHLSRPRS” (Fig 5B).

SIRT5 had one MTS sequence “MRPLL IAPGR FISQL CCRRK PPASP QSKIC” in the NH-terminus of the protein (Fig 5C).

### The existence of SRF regulatory sequences in the promoter region of siruin genes

SRF has been reported as an upstream regulator of the human SIRT2 gene, which contains a serum response element (SRE, or classic CArG motif) that is bound to SRF [24]. To examine whether the other sirtuin gene promoters may also contain classic CArG motif or CArG-like motif, the promoter regions of all the seven mouse sirtuin genes were analyzed using the promoter analysis software tools (Methods and materials). As show in Table-2, there were CArG-like sequences in the −5 kb promoter region of all seven mouse sirtuin genes. In addition, there was a classic CArG motif “CCAAAAAAGG” in the SIRT5 gene promoter. These data indicate that SRF and SRF cofactors may partly contribute to the activity of sirtuin genes via transcriptional regulation.

**Table 2.** SRF binding site (CArG-like and classic CArG motifs) in mouse sirtuin gene promoters.

| Promoter region (-5 kb) | Sequence | Motif |
| --- | --- | --- |
| SIRT1 Promoter | CCTGTTATGC | CArG-like |
|  | ACACAAAAGG | CArG-like |
| SIRT2 Promoter | CCAAATTTGT | CArG-like |
| SIRT3 Promoter | CCTATATATG | CArG-like |
|  | CCATCATATG | CArG-like |
|  | CCTAATGTGG | CArG-like |
| SIRT4 Promoter | TCCAATTTGG | CArG-like |
| SIRT5 Promoter | CCAAAAAAGG | classic CArG |
|  | CCAAATTGGA | CArG-like |
|  | CCAAAAAGGG | CArG-like |
|  | ACCTATTTGG | CArG-like |
| SIRT6 Promoter | CCGAATTTAG | CArG-like |
| SIRT7 Promoter | CCTCTGTATG | CArG-like |
|  | CACATTATGG | CArG-like |
|  | CCAAATACGC | CArG-like |

### Comparison of exon number of sirtuin genes between mouse and human

It has been reported that constitutive splicing events have a high level of conservation, the alternative splicing junctions usually bear a lower level of conservation across species [33,34]. For instance, the mouse SIRT1v2 isoform retained the NLS during alternatice splicing, whereas the similar SIRT1 isoform in human has not been reported [10].

Examination of the genomic sequences and mRNA sequences showed the difference of exon numbers between human and mouse sirtuin gene family. Six sirtuin genes had different exon numbers in human versus mouse (Fig 6A). The human SIRT1 gene had two more exons than mouse SIRT1 genes (11 vs 9 exons); the human SIRT5 gene had four more exons than mouse SIRT5 gene (17 vs 13 exons).

**Fig 6.** The divergence of sirtuin gene sequences between human and mouse. A. The table showed the exon number difference between human sirtuin and mouse genes. Only SIRT2 gene had the same exon number in human and mouse genome. B-D. Comparison of exon and splicing events between human and mouse SIRT2 isoforms. C. Both human and mouse SIRT2v1 and v2 had 16 exons. D. Both SIRT2v1 and v2 had 15 exons, in which the exon-2 was spliced out. E. Three exons skipped in both human and mouse SIRT2v3. The ex1 (exon 1), ex13, ex14 skipped in human SIRT2v3, while Ex2, ex3, and ex4 skipped in mouse SIRT2v3. E. Comparison of SIRT2 non-coding region (promoter region) vs mRNA. There is 86% homology in mRNA between human and mouse, but 35-78% homology in promoter region between human and mouse.

The SIRT2 gene was the only gene in the sirtuin family that had same number of exons in both human and mouse (17 vs 17 exons) (Fig 6A). There were 3 SIRT2 isoforms in both human and mouse. Both the human and mouse SIRT2v1 and v2 had the same number of exons (Figs 6B and 6C), but the different exons were skipped in SIRT2v3 in human versus mouse (Fig 6D). Further analysis showed that there was 86% homology in the mRNA sequences, but lower homology in the non-coding promoter regions between human and mouse SIRT2 genes, ranging from 35% to 78% (Fig 6E).

### The expression of sirtuin genes in response to serum deprivation and restoration

Since CArG-like and/or classic CArG sequences were found in the promoter regions of all seven sirtuin genes, the expression of sirtuin isoforms in response to the serum stresses were examined in a cellular stress model with serum deprivation and restoration, which reflected the blood supply and nutrient level changes during the ischemia and reperfusion conditions.

Most of the sirtuin isoforms were upregulated at approximately 6 hours after serum deprivation, including SIRT1v1,SIRT2v1, v2, v3, SIRT3, SIRT4, and SIRT5. However, SIRT1v2 was down-regulated. SIRT6 was up-regulated at a high level at 24 hours, and SIRT7 was increased at 18 hours after serum deprivation (Fig 7A).

**Fig 7.** Sirtuin gene and isoform expression in response to serum deprivation and restoration. A. Sirtuin gene and isoform expression in response to serum deprivation. The gene expression was measured after 3 hours (3h), 6h, 18h, 24h and 48h of serum deprivation. B. Sirtuin gene and isoform expression in response to serum deprivation. The gene expression was measured after 3 hours (3h), 6h, 18h, 24h and 48h of serum restoration.

Most sirtuin isoform expression levels were up-regulated in response to serum restoration after deprivation, where the levels reached the peak at around 48 hours after serum restoration (Fig 7B).

## Discussion

The present study has several major findings. A total of 15 sirtuin isoforms were identified in the mouse genome. Among the seven sirtuin genes (SIRT1-SIRT7), six sirtuin genes had two or more isoforms, whereas SIRT5 gene had one isoform. Alternative splicing increased sirtuin transcriptomic and proteomic diversity, and generated the sirtuin variants, most of which lacked part of the coding region sequence. The changes in the sirtuin coding region were predicted to cause protein structural change. In addition, the NLS, NoLS, and MTS sequences in the sirtuin isoforms were analyzed, and found to predict the intracellular localization of each sirtuin isoform. Analysis of the sirtuin gene promoter regions showed both classic CArG motif and CArG-like motif in all seven sirtuin genes, indicating that a potential regulatory mechanism may be associated with SRF and its cofactors.

Thanks to the advancement of large-scale transcriptomic and proteomic analyses, it is now appreciated that alternative splicing occurs in transcripts produced by the majority of protein coding genes containing multiple exons [6,28]. Indeed, the seven mouse sirtuin genes have multiple exons, ranging from 7 exons (i.e SIRT4) to 17 exons (i.e. SIRT2), and most of them have more than one isoform. Since exon skipping is a common event during splicing, the alternatively spliced isoforms are usually shorter than the constitutively spliced main isoform (v1 isoform). Compared to the v1 isoform, most v2 and v3 isoforms have lost one or more exons during the pre-mRNA splicing. Most exon skipping events occur near the N-terminus, but the SIRT7v2 isoform lost an exon in the C-terminus.

The human and mouse genomes share similar long-range sequence organization, and most of the coding sequences are highly conserved and homologous. The intron sequences tend to have lower homology among species [10,29]. Both human and mouse genomes contain seven sirtuin genes. Even though the coding regions and protein sequences of the sirtuin genes are highly homologous between human and mouse, the genomic sequences in the non-coding DNA regions are likely to be of at a lower conservative level. This lower conservation in non-coding DNA sequences could be found in the exon and coding region distribution in terms of exon number. Among the seven sirtuin genes, six sirtuin genes contained different numbers of exons between human versus mouse, while only the sirtuin-2 gene contained 17 exons in both the human and mouse genomes.

The broad divergence in genomic sequences may result in different sirtuin isoforms in human versus mouse. Additionally, many alternative splicing events and isoforms could be species-specific. Recently, we reported the identification of 25 human sirtuin isoforms, many of them have not been found in the mouse [33]. We have previously found a classic CArG motif (CCATAATAGG) in the promoter region of the human sirtuin-2 gene. We have observed that SRF binds tightly to the SIRT-2 gene promoter in the electrophoretic mobility shift assay (EMSA); and SIRT2 gene expression is repressed by a Rho/SRF inhibitor in response to serum stimulation, indicating that SIRT2 gene is a downstream target of the Rho/SRF signaling pathway [24].

In the present study, the classic CArG and CArG-like motifs were also found in the proximal promoter regions of the mouse sirtuin genes. These data further indicate that the some sirtuin genes are likely regulated by the signaling pathways associated with SRF, likely under certain physiological and pathological conditions, including serum restoration following serum deprivation [30]. The activation of SRF subsequently increases SRF binding activity to its cognate binding sequence, thereby activating or repressing its target genes [31]. In the present study, we utilized the cellular stress model of “serum deprivation and restoration” to mimic the blood supply and nutrient level changes in the pathological condition of “ischemia and reperfusion”, which is characterized by an initial restriction of blood supply to an organ followed by the subsequent restoration of blood perfusion [32], which may happen in various cardiac and vascular diseases, as well as organ transplantation and cardiothoracic, vascular and general surgery.

The sirtuin genes are considered to be nutrient sensors, and sirtuin catalytic activity is regulated in response to a change of nutritional status, especially, the intracellular level of NAD^+^ [33]. Sirtuin genes are also regulated by transcriptional regulation. For instance, the transcriptional factors, early growth response 2, 3 (EGR2, EGR3), bind to sirtuin-1 gene promoter thereby regulate the sirtuin-1 gene expression [34]. Additionally, PPARβ/δ regulates the human SIRT1 gene transcription via Sp1 [35]. The human SIRT2 gene contains a classic CarG motif in its promoter, and is regulated by SRF and Rho/SRF signaling pathway. Analysis of the promoter sequences suggested that multiple transcriptional regulatory sites exist in sirtuin genes, including the classic CArG, CArG-like motifs, EGR1, EGR2, EGR3 and Sp1. Therefore, we propose a potential transcriptional regulatory mechanism of sirtuin genes involving SRF and other transcription factors/cofactors (Fig 8).

**Fig 8.**
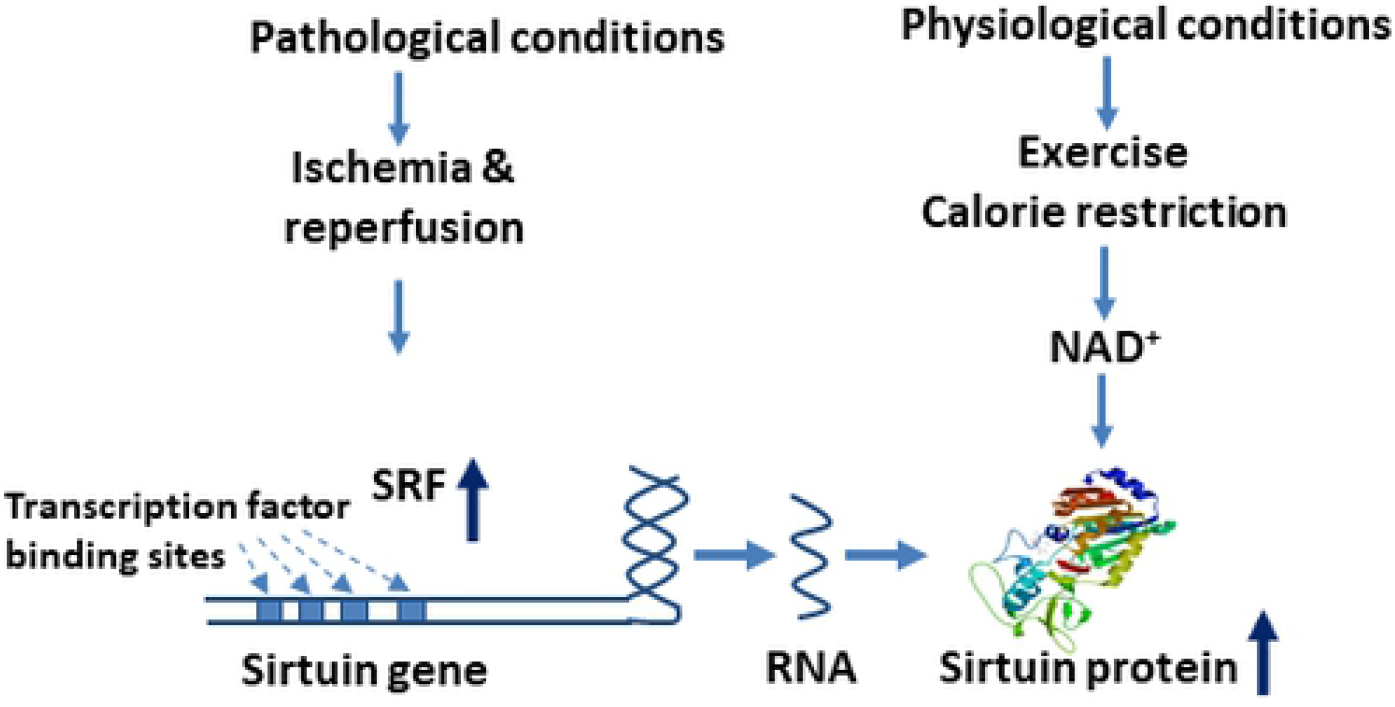
A schematic diagram of sirtuin gene regulation under physiological and pathological conditions. We propose a potential model of transcriptional regulation of sirtuin genes by SRF in pathological conditions, including ischemia & reperfusion. 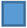 Transcription factor binding sites in sirtuin gene promoter, including EGR2, EGR3, Sp1, CArG and CArG-like, etc.

**<u>In conclusion</u>,** it is possible that SRF may play an important role in the transcriptional regulation of sirtuin genes, at least the human SIRT2 and mouse SIRT5 genes that contain the classic CArG motif, in ischemia and reperfusion, a condition where SRF is likely to be strongly activated.

## Acknowledgments

We thank Shakshi Sharma, and Yingni Che for technical assistance.

## Ethical Approval

The studies were conducted with Institutional Review Board approval from the University of Arkansas for Medical Sciences, in accordance with the NIH Guiding Principles for Research Involving Animals and Human Beings.

## Data Availability Statement

All the mRNA, protein and genomic DNA sequences of the sirtuin genes are available at The National Center for Biotechnology Information (NCBI) database. Additional data are available upon request.

## Consent for publication

All authors have read and agreed to the published version of the manuscript.

## Authors’ Contributions

Conceived and designed the experiments: X.Z., J.Y.W. Performed the experiments: X.Z., F.S.A., J.C.. Analyzed the data: X.Z., F.S.A., G.A., J.Y.W. Contributed reagents/materials/analysis tools: G.A. Wrote the paper: X.Z., J.Y.W. Statistical analysis: X.Z., G.A., F.S.A.. All authors have read and agreed to the published version of the manuscript.

## References

1. Han H, Braunschweig U, Gonatopoulos-Pournatzis T, Weatheritt RJ, Hirsch CL, et al. (2017) Multilayered Control of Alternative Splicing Regulatory Networks by Transcription Factors. Mol Cell 65: 539–553 e537.

2. Irimia M, Blencowe BJ (2012) Alternative splicing: decoding an expansive regulatory layer. Curr Opin Cell Biol 24: 323–332.

3. Licatalosi DD, Darnell RB (2010) RNA processing and its regulation: global insights into biological networks. Nat Rev Genet 11: 75–87.

4. Nilsen TW, Graveley BR (2010) Expansion of the eukaryotic proteome by alternative splicing. Nature 463: 457–463.

5. Pan Q, Shai O, Lee LJ, Frey BJ, Blencowe BJ (2008) Deep surveying of alternative splicing complexity in the human transcriptome by high-throughput sequencing. Nat Genet 40: 1413–1415.

6. Wang ET, Sandberg R, Luo S, Khrebtukova I, Zhang L, et al. (2008) Alternative isoform regulation in human tissue transcriptomes. Nature 456: 470–476.

7. Frye RA (2000) Phylogenetic classification of prokaryotic and eukaryotic Sir2-like proteins. Biochem Biophys Res Commun 273: 793–798.

8. Thompson M, Bixby R, Dalton R, Vandenburg A, Calarco JA, et al. (2019) Splicing in a single neuron is coordinately controlled by RNA binding proteins and transcription factors. Elife 8.

9. Cheng C, Yaffe MB, Sharp PA (2006) A positive feedback loop couples Ras activation and CD44 alternative splicing. Genes Dev 20: 1715–1720.

10. Zhang X, Ameer FS, Azhar G, Wei JY (2021) Alternative Splicing Increases Sirtuin Gene Family Diversity and Modulates Their Subcellular Localization and Function. Int J Mol Sci 22.

11. O’Leary NA, Wright MW, Brister JR, Ciufo S, Haddad D, et al. (2016) Reference sequence (RefSeq) database at NCBI: current status, taxonomic expansion, and functional annotation. Nucleic Acids Res 44: D733–745.

12. Chang HC, Guarente L (2014) SIRT1 and other sirtuins in metabolism. Trends Endocrinol Metab 25: 138–145.

13. Corpet F (1988) Multiple sequence alignment with hierarchical clustering. Nucleic Acids Res 16: 10881–10890.

14. Sigrist CJ, de Castro E, Cerutti L, Cuche BA, Hulo N, et al. (2013) New and continuing developments at PROSITE. Nucleic Acids Res 41: D344–347.

15. Waterhouse A, Bertoni M, Bienert S, Studer G, Tauriello G, et al. (2018) SWISS-MODEL: homology modelling of protein structures and complexes. Nucleic Acids Res 46: W296–W303.

16. Kosugi S, Hasebe M, Tomita M, Yanagawa H (2009) Systematic identification of cell cycle-dependent yeast nucleocytoplasmic shuttling proteins by prediction of composite motifs. Proc Natl Acad Sci U S A 106: 10171–10176.

17. Nair R, Carter P, Rost B (2003) NLSdb: database of nuclear localization signals. Nucleic Acids Res 31: 397–399.

18. la Cour T, Kiemer L, Molgaard A, Gupta R, Skriver K, et al. (2004) Analysis and prediction of leucine-rich nuclear export signals. Protein Eng Des Sel 17: 527–536.

19. Xu D, Marquis K, Pei J, Fu SC, Cagatay T, et al. (2015) LocNES: a computational tool for locating classical NESs in CRM1 cargo proteins. Bioinformatics 31: 1357–1365.

20. Brameier M, Krings A, MacCallum RM (2007) NucPred--predicting nuclear localization of proteins. Bioinformatics 23: 1159–1160.

21. Claros MG, Vincens P (1996) Computational method to predict mitochondrially imported proteins and their targeting sequences. Eur J Biochem 241: 779–786.

22. Emanuelsson O, Nielsen H, Brunak S, von Heijne G (2000) Predicting subcellular localization of proteins based on their N-terminal amino acid sequence. J Mol Biol 300: 1005–1016.

23. Fukasawa Y, Tsuji J, Fu SC, Tomii K, Horton P, et al. (2015) MitoFates: improved prediction of mitochondrial targeting sequences and their cleavage sites. Mol Cell Proteomics 14: 1113–1126.

24. Zhang X, Azhar G, Wei JY (2017) SIRT2 gene has a classic SRE element, is a downstream target of serum response factor and is likely activated during serum stimulation. PLoS One 12: e0190011.

25. Zhang X, Azhar G, Wei JY (2012) The expression of microRNA and microRNA clusters in the aging heart. PLoS One 7: e34688.

26. Schmittgen TD, Livak KJ (2008) Analyzing real-time PCR data by the comparative C(T) method. Nat Protoc 3: 1101–1108.

27. Scott MS, Troshin PV, Barton GJ (2011) NoD: a Nucleolar localization sequence detector for eukaryotic and viral proteins. BMC Bioinformatics 12: 317.

28. Kelemen O, Convertini P, Zhang Z, Wen Y, Shen M, et al. (2013) Function of alternative splicing. Gene 514: 1–30.

29. Thanaraj TA, Clark F, Muilu J (2003) Conservation of human alternative splice events in mouse. Nucleic Acids Res 31: 2544–2552.

30. Norman C, Runswick M, Pollock R, Treisman R (1988) Isolation and properties of cDNA clones encoding SRF, a transcription factor that binds to the c-fos serum response element. Cell 55: 989–1003.

31. Miano JM, Long X, Fujiwara K (2007) Serum response factor: master regulator of the actin cytoskeleton and contractile apparatus. Am J Physiol Cell Physiol 292: C70–81.

32. Eltzschig HK, Eckle T (2011) Ischemia and reperfusion--from mechanism to translation. Nat Med 17: 1391–1401.

33. Bonkowski MS, Sinclair DA (2016) Slowing ageing by design: the rise of NAD(+) and sirtuin-activating compounds. Nat Rev Mol Cell Biol 17: 679–690.

34. Gao B, Kong Q, Kemp K, Zhao YS, Fang D (2012) Analysis of sirtuin 1 expression reveals a molecular explanation of IL-2-mediated reversal of T-cell tolerance. Proc Natl Acad Sci U S A 109: 899–904.

35. Okazaki M, Iwasaki Y, Nishiyama M, Taguchi T, Tsugita M, et al. (2010) PPARbeta/delta regulates the human SIRT1 gene transcription via Sp1. Endocr J 57: 403–413.

